# Diffusion MRI Head Motion Correction Methods are Highly Accurate but Impacted by Denoising and Sampling Scheme

**DOI:** 10.1101/2022.07.21.500865

**Authors:** Matthew Cieslak, Philip A. Cook, Tinashe M. Tapera, Hamsanandini Radhakrishnan, Mark Elliott, David R. Roalf, Desmond J. Oathes, Dani S. Bassett, M. Dylan Tisdall, Ariel Rokem, Scott T. Grafton, Theodore D. Satterthwaite

## Abstract

Correcting head motion artifacts in diffusion-weighted MRI (dMRI) scans is particularly challenging due to the dramatic changes in image contrast at different gradient strengths and directions. Head motion correction is typically performed using a Gaussian Process model implemented in FSL’s Eddy. Recently, the 3dSHORE-based SHORELine method was introduced to correct any dMRI sequence that has more than one shell. Here we perform a comprehensive evaluation of both methods on realistic simulations of a software fiber phantom that provides known ground-truth head motion. We demonstrate that both methods perform remarkably well, but that performance can be impacted by sampling scheme, the pervasiveness of head motion, and the denoising strategy applied before head motion correction. Our study also provides an open and fully-reproducible workflow that could be used to accelerate evaluation studies of other dMRI processing methods in the future.

**HIGHLIGHTS:**

- Both Eddy and SHORELine head motion correction methods performed quite well on a large variety of simulated data
- Denoising with MP-PCA can improve head motion correction performance when Eddy is used
- SHORELine effectively corrects motion in non-shelled acquisitions

## INTRODUCTION

Diffusion-weighted MRI (dMRI) modulates the MR signal to encode information about the distribution of water diffusion, which is constrained by the orientation and permeability of tissue (Basser, 1995; Callaghan, 1993; Stejskal & Tanner, 1965). This method has become widely used to non-invasively image the structural properties of white matter in the brain. Over the course of three decades, dMRI sequences have advanced to measure signal in many directions (e.g., higher angular resolution) and diffusion sensitizations (Tuch, 2004; Wedeen et al., 2005) with most modern sequences capturing hundreds of images over the course of 10 to 30 minutes of scanning.

Any scanning sequence where multiple images are acquired over time is highly susceptible to artifacts related to head motion during the scan. The effects of head motion during functional MRI (fMRI), another imaging technique that acquires images in a series over time, are well-known and typically addressed by simply aligning each image to a reference image using a rigid or affine transformation (Jenkinson et al., 2002), followed by further corrections to the time series data in each voxel (Ciric et al., 2018). The use of a single reference image works well for spatially correcting fMRI because the contrast and SNR remain relatively constant over the acquisition.

In contrast to fMRI, dMRI sequences acquire images that can have dramatically different spatial contrasts and SNR depending on the diffusion-encoding gradient moment (i.e., the *b*-value in s/mm^2^) and direction. The set of directions and *b*-values that define a dMRI sampling scheme are what allow the method to estimate the ensemble average diffusion propagator (EAP) in each voxel (Callaghan, 1993). However, such differences in contrast also preclude the use of a single image as the registration target for head motion correction. Instead, for each *b*>0 image in the dMRI series, an image with similar spatial contrast must be generated as if it were aligned with all other images in the dMRI series. Each image can then be registered to the target image, thereby correcting the effect of bulk head motion in each volume.

At present the most widely-used method for dMRI head motion correction is Eddy (Andersson & Sotiropoulos, 2015), which is included in the fMRIB software library (FSL). Eddy has been broadly adopted, including by large imaging consortia such as the Human Connectome Project (Glasser et al., 2013), the UK BioBank (Alfaro-Almagro et al., 2018), and a version of the Healthy Brain Network (Richie-Halford et al., 2022). In addition to estimating bulk head motion, Eddy estimates and corrects spatial warping related to eddy currents, fills in dropped slices (Andersson et al., 2016), estimates intra-volume motion, and optionally incorporates susceptibility distortion correction if a fieldmap is estimated using the TOPUP tool (Andersson & Sotiropoulos, 2016). Many of these features rely on Eddy’s algorithm for generating registration targets. Eddy operates on *shelled* dMRI sequences, which acquire multiple gradient directions at the same *b*>0 value. Given that all images on the same shell are sampling the surface of a sphere in *q*-space, the differences in their signal can be represented as a Gaussian process (GP) on the azimuth and elevation coordinates on the unit sphere *S*^2^. Eddy estimates one GP per shell and uses the GP to produce registration targets. A rigid registration is performed between each *b*>0 image and its GP prediction, followed by a linear or quadratic warp in the Phase Encoding Direction of the dMRI acquisition to correct for distortions related to eddy currents.

Although shelled acquisitions are popular, there are other methods of sampling *q*-space that have unique advantages. Diffusion Spectrum Imaging (DSI) samples a Cartesian grid in *q-* space, enabling the direct reconstruction of the EAP with a simple Fourier transform (Wedeen et al., 2005). Shelled schemes require more complex modeling and leave open a question with no universal answer: at which *b*-values should one acquire shells? DSI scans typically have required the acquisition of more than 200 images, resulting in long scan times—particularly when multiband imaging is not available. However, sparse, random subsets of the Cartesian grid scheme along with a compressed-sensing reconstruction approach (CS-DSI) have been shown to provide comparable EAP reconstructions to the full grid sampling scheme at a fraction of the scanning time (Merlet & Deriche, 2013; Paquette et al., 2015). Yet, there is no widely accepted head motion correction for these non-shelled schemes, which limits the application of DSI in translational research.

Recently the SHORELine algorithm (Cieslak et al., 2021) was introduced as a method to generate registration targets for any dMRI sequence with both radial and angular variability in its *q*-space sampling scheme. SHORELine is a cross-validated method where, for each *b*>0 image, the 3dSHORE basis (Özarslan et al., 2013) set is fit to all other images using L2-regularization. A registration target for the left-out image is estimated from the 3dSHORE fit, and the image is registered with a Rigid (6DOF) or optional Affine (12DOF) transform using ANTs **(Figure 1)**. No eddy current correction is explicitly attempted. This process is repeated up to 2 times based upon user specifications. The 3dSHORE basis functions are defined in three-dimensional space (*R*^3^) and are therefore appropriate for multi-shelled, Cartesian and sparse/random sampling schemes. Like Eddy, SHORELine can incorporate a susceptibility distortion correction along with head motion correction in a single interpolation. The original evaluation study showed an overall improvement in the Neighboring DWI Correlation (NDC) quality measure of non-shelled sampling schemes compared to the unprocessed data (Cieslak et al., 2021).

**Figure 1:**
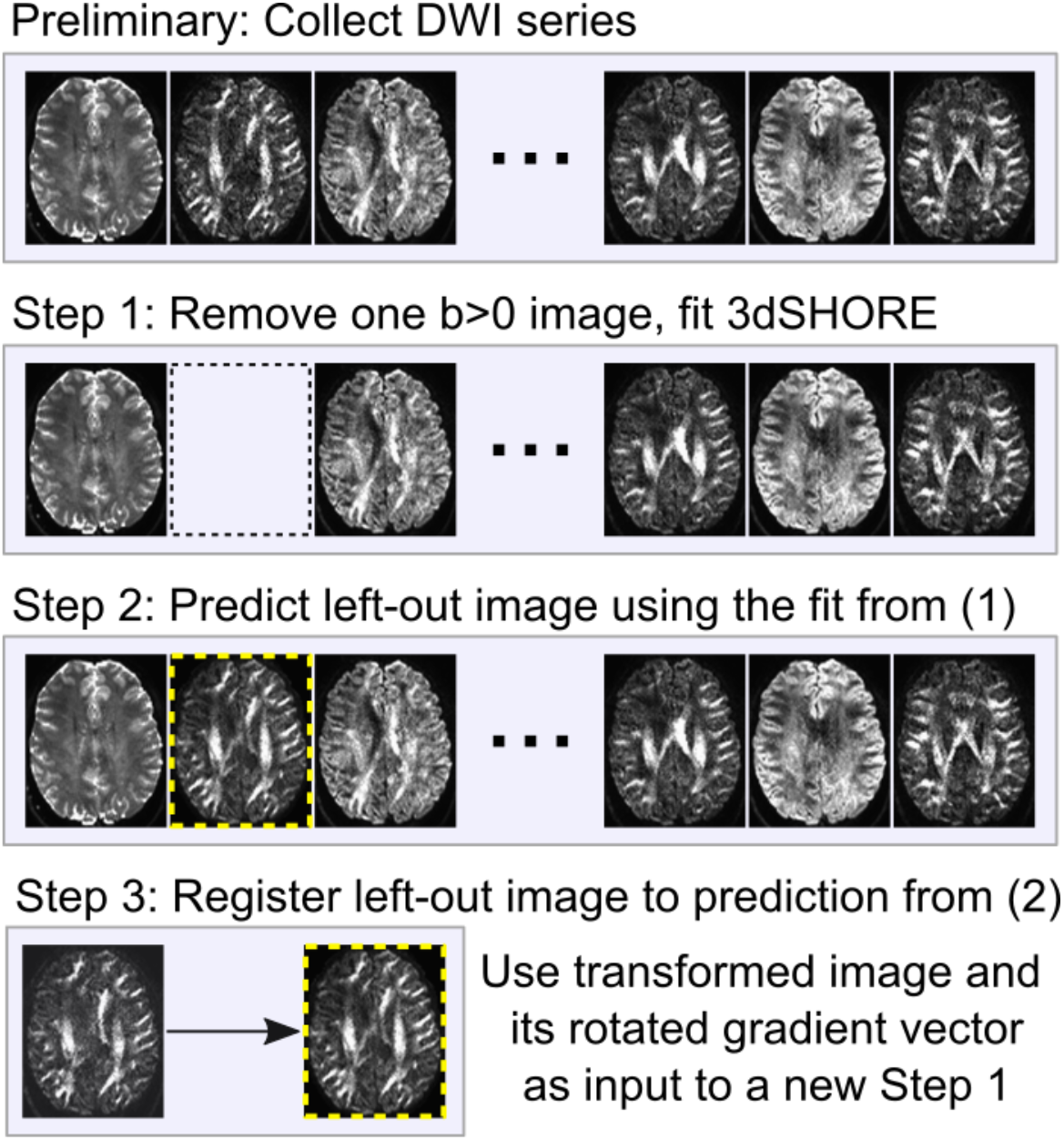
SHORELine. Images shown here are from an actual scan using the CS-DSI sampling scheme. The slice in the yellow-dashed box is the SHORELine-predicted slice for the left-out volume. The actual left-out slice is shown on the left in Step 3. The contrast in the predicted slice is visually very similar to the actual left-out slice, which enables standard image registration methods to work.

Independent of head motion correction, denoising algorithms are often applied prior to head motion correction. The application of MP-PCA denoising (Veraart et al., 2016) is enabled by default in both QSIPrep (Cieslak et al., 2021) and the `MRtrix3_connectome` (Smith & Connelly, 2020) pipelines. However, denoising is not applied in the HCP (Glasser et al., 2013) or UK Biobank (Alfaro-Almagro et al., 2018) pipelines. There are reasonable arguments both for and against performing denoising prior to motion correction. The registration targets generated by Eddy and SHOREline may be more realistic if their underlying models are fit with less-noisy data. However, denoising is not recommended by the developers of Eddy because it will affect how noise is distributed in the dMRI signal, thereby possibly violating assumptions of the GP (Andersson, 2019).

Both Eddy and SHORELine are considerably more complex than standard fMRI head motion correction methods. However, there has not been a systematic evaluation of how well these methods perform on commonly acquired sequences with different levels of head motion, nor has there been a systematic evaluation on the effect of denoising dMRI data prior to head motion correction. To address this gap, here we simulated hundreds of thousands of dMRI images from common shelled and non-shelled schemes incorporating known head motion and realistic MR artifacts to determine how accurate these methods are in estimating true head motion with and without denoising.

## METHODS

### Simulation of images and motion

MITK FiberFox (Neher et al., 2014) was used to simulate the entire dMRI series for a set of commonly acquired sampling schemes, including ABCD, HCP, a Cartesian DSI half sphere and a Cartesian/random CS-DSI. The simulation used streamline segments from the ISMRM 2015 fiber phantom (http://www.tractometer.org/ismrm_2015_challenge/data) to define a fiber ODF in each voxel and convert it into a diffusion ODF to generate an MRI signal. The MRI signal was combined with eddy currents, thermal noise, and susceptibility distortion artifacts. Simulation parameters used here are identical to the ISMRM simulated phantom, except that the original study contained only *three volumes* with simulated head motion, whereas we simulated hundreds of thousands of volumes under different sampling schemes and conditions (see below). A docker image of the exact version of Fiberfox can be downloaded at https://hub.docker.com/r/pennbbl/fiberfox using tag 1.0.

To introduce controlled head motion, a random rigid transform was applied to the streamline data before the MR signal was simulated (Hering et al., 2014). Both translation and rotation about each axis were sampled from a random uniform distribution with a maximum absolute displacement of 5 mm and a maximum absolute rotation of 5 degrees **(Figure 2)**. Individual dMRI series were created by replacing a random subset of images from the no-motion simulation with the motion-included simulated images. Unique dMRI scans were generated to have a specific *motion prevalence* such that 15%, 30%, or 50% of the volumes in the series included head motion. A total of 30 unique scans were generated for each sampling scheme for each of the three motion prevalence values, yielding 90 simulated complete series per scheme. In total this process resulted in 360 unique simulated dMRI scans.

**Figure 2:**
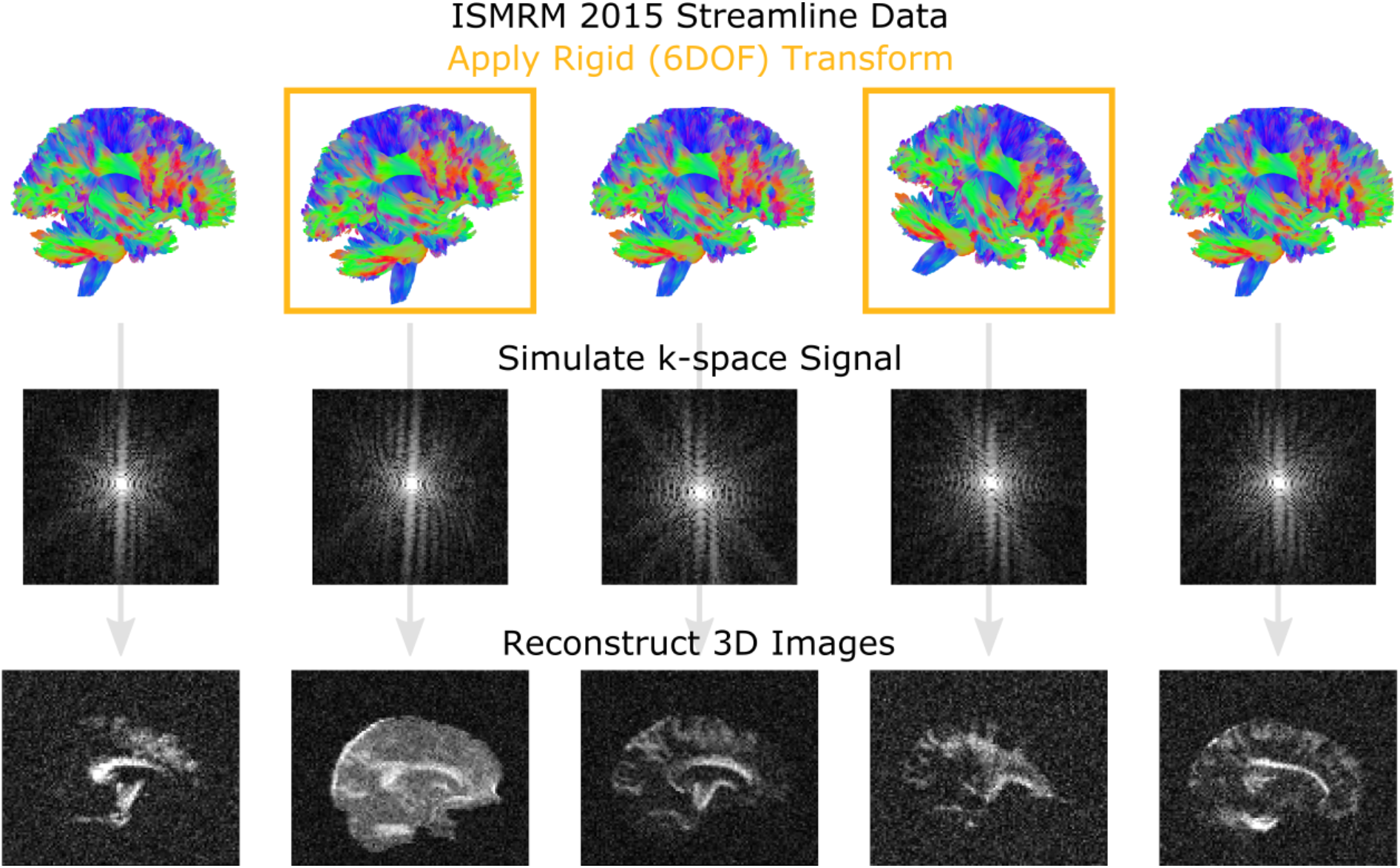
Simulation of head motion in DWI images with Fiberfox. Fiberfox simulates MRI data from a set of streamlines. Here we show the streamlines used for benchmarking (top row). We introduced motion to our test data by applying rigid (6-DOF) transformations to the streamlines before the volumes were simulated. The diffusion restriction introduced by the fibers represented by the streamlines is included in the signal attenuation in each voxel, which is then “acquired” in *k*-space where artifacts can be introduced (middle row). Finally, the *k*-space data is reconstructed into realistic 3D volumes that are used for benchmarking (bottom row).

### Image processing

Eddy and SHORELine were run on each simulated scan using QSIPrep (v0.14.3). This version of QSIPrep included FSL version 6.0.3. Eddy was run twice, once with Linear and once with Quadratic models. However, a comparison of the two approaches revealed that these options do not appear to have a substantial impact on head motion estimation. Accordingly, we report the results from the Eddy with the quadratic setting because this is the way it is typically used as part of the HCP Pipelines. SHORELine was also run twice, once with a Rigid (6DOF) and once with an Affine (12DOF) transformation model. The rigid model had minor benefits and also a shorter run time; as such, the results for the rigid model are used to characterize SHORELine’s performance. Each configuration was run with and without MP-PCA denoising (from MRTrix 3.0.3) prior to head motion correction. In all, a total of 2,160 QSIPrep preprocessing runs were executed, encompassing the processing of 375,840 *b*>0 images (**Table 1**).

**Table 1.**
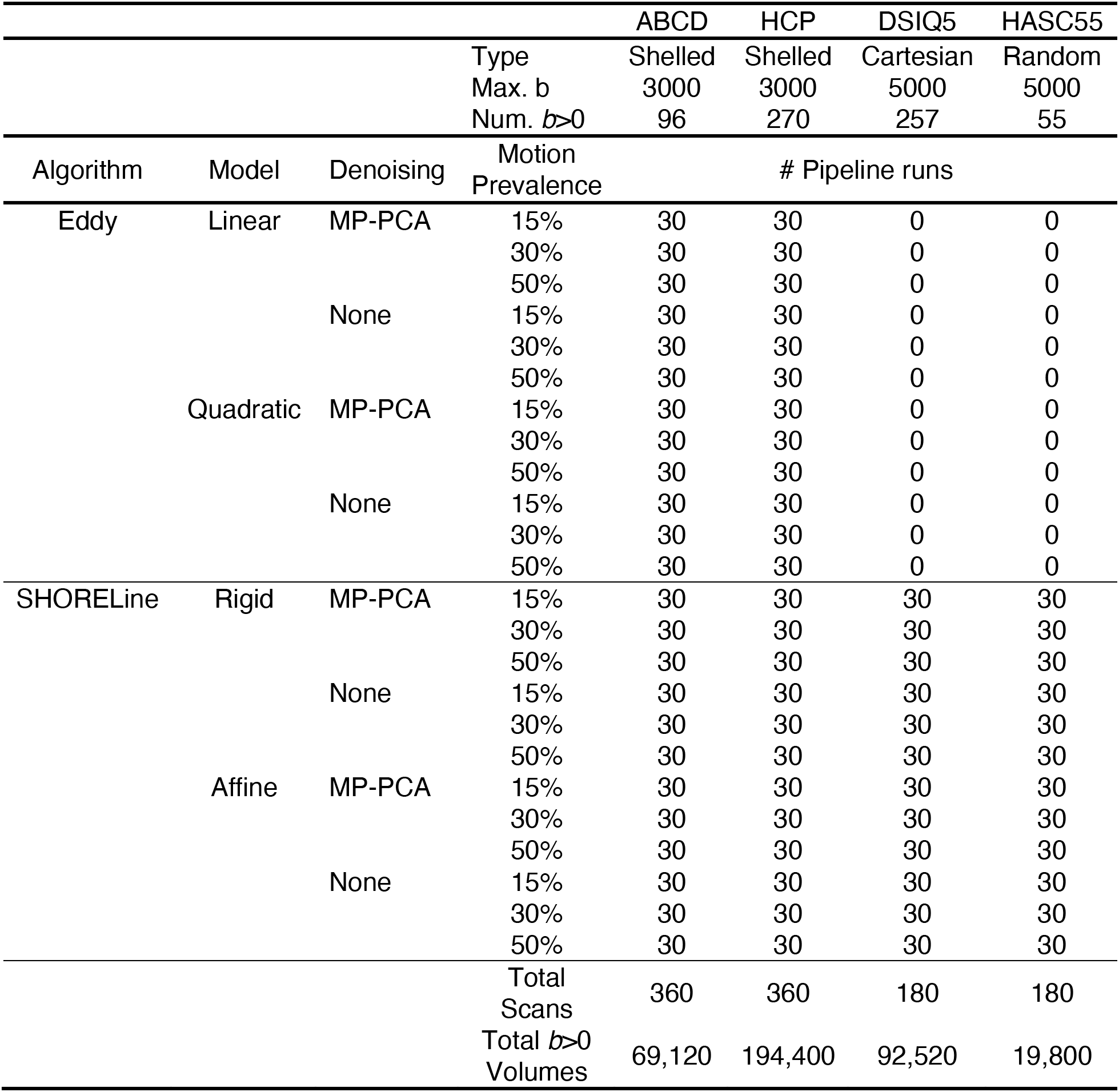
Properties of simulated datasets and how they were processed. Each sequence was simulated such that 15%, 30%, and 50% of volumes contained head motion. Each level of motion was processed both with and without denoising.

Importantly, to ensure reproducibility, the entire benchmarking experiment was run using the FAIRly big workflow (Wagner et al., 2022). This approach uses DataLad to track and distribute the data and code used during data analysis. The entire FiberFox phantom dataset along with a Singularity image of the software used to run the processing are publicly available. Each run of QSIPrep was recorded as a git commit and can be reproduced locally by anyone with DataLad who clones from the repository (https://github.com/PennLINC/dMRI_HMC_Benchmark).

### Outcome measures

Performance was evaluated according to multiple metrics. First, the mean error in head motion parameter estimation was calculated to determine whether the algorithm is an unbiased estimator. Second, to characterize the expected error of the estimators, we calculated the RMSE of the estimated head motion. Low RMSE reflects accurate head motion estimation and higher RMSE indicates greater estimation error. To understand the factors that affect motion parameter estimation, linear models were fit with root mean squared error (RMSE) as the dependent variable. As rotation and translation are in different units, we fit two linear models of RMSE for translation and rotation separately. The relative performance between SHORELine and Eddy was calculated by subtracting SHORELine’s RMSE from Eddy’s, resulting in positive values when SHORELine’s was more accurate at estimating motion parameters than Eddy.

Third, we compared the interpolation-related smoothness of the corrected images. Image smoothness is a measure of blurring during preprocessing, which reduces anatomical detail; preprocessing should seek to minimize the introduction of additional image smoothness. We estimated the full-width at half-maximum (FWHM) of the mean *b*=0 image in the preprocessed data. Fourth, we evaluated a summary measure of data quality—the neighboring DWI Correlation (NDC) (Yeh et al., 2019). NDC summarizes the pairwise spatial correlation between each pair of dMRI volumes that sample the closest points in *q*-space; lower values reflect reduced data quality, driven by noise and misalignment between dMRI volumes.

## RESULTS

### Both Eddy and SHORELine accurately correct simulated head motion

Both Eddy and SHORELine demonstrated excellent performance in correcting head motion. While collapsing across all experimental conditions, the *mean error* (calculated as the mean difference between the estimated motion parameter and the ground truth motion parameter) in estimated head rotation was very small: only 0.194 degrees. Similarly, the mean error in estimated head translation was only ~1/100^th^ of a voxel: 0.012 mm. Such miniscule mean errors suggest that both Eddy and SHORELine are accurate and unbiased estimators of head motion parameters. Mean errors and RMSE (the first and second moments of the error distribution) are provided in **Table 2**.

**Table 2.** Mean error of head motion correction methods

| Sampling Scheme | Method | *Translation (mm)* |  | *Rotation (°)* |  |
| --- | --- | --- | --- | --- | --- |
|  |  | Mean Error | RMSE | Mean Error | RMSE |
| ABCD | Eddy + MP-PCA | 0.016 | 0.71 | 0.096 | 0.70 |
|  | Eddy | 0.071 | 1.1 | 0.27 | 2.3 |
|  | SHORELine | −0.086 | 0.80 | −0.037 | 1.6 |
| HCP | Eddy + MP-PCA | −0.044 | 0.73 | 0.0093 | 0.56 |
|  | Eddy | 0.063 | 0.71 | −0.063 | 1.0 |
|  | SHORELine | −0.051 | 0.56 | 0.48 | 0.86 |
| DSIQ5 | SHORELine | 0.10 | 0.61 | 0.24 | 0.96 |
| HASC55 | SHORELine | 0.059 | 0.99 | 0.33 | 1.5 |

### Head motion correction accuracy varies by sampling scheme and denoising

Next, we evaluated what factors impacted error (hereafter referring to RMSE) following head motion correction (**Figure 3**). As described below, results indicate that error following motion correction is primarily due to uncontrolled factors such as the amount of head motion present in the data but also preprocessing choices (use of denoising) and factors related to experimental design (sampling scheme); see **Supplementary Table 1** for complete statistical results.

**Figure 3:**
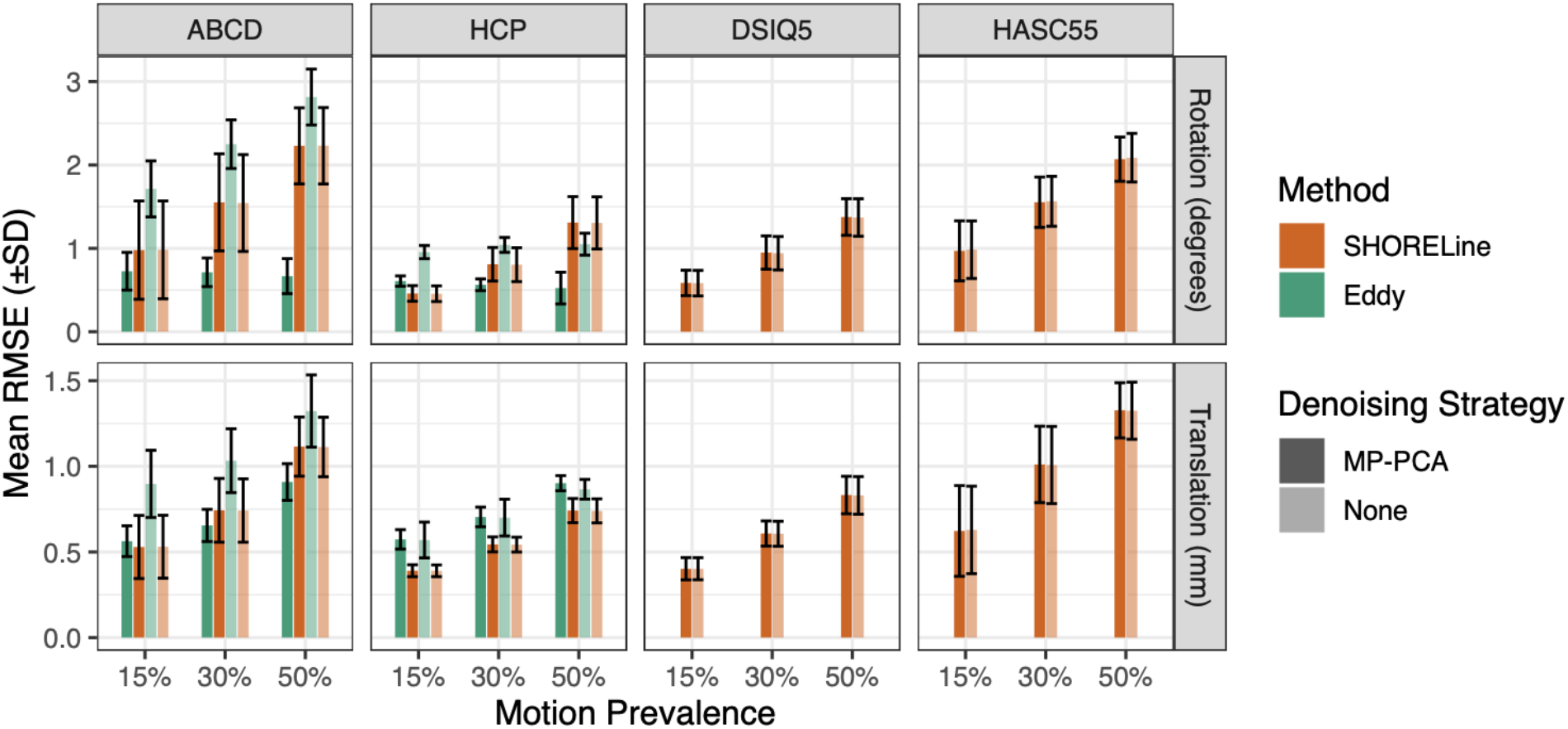
Head motion correction error varies by motion present in input data, sampling scheme, and de-noising. Means and standard deviations of the motion parameter estimate RMSE are plotted for both Eddy and SHORELine. Eddy does not support non-shelled schemes and therefore could not be evaluated for DSIQ5 and HASC55. Error bars reflect the standard deviation of the RMSE for the sample, each of which consists of 30 simulated scans. Error varied by sampling scheme, with lower errors present in simulated HCP data than ABCD data. Notably, greater motion in the input data was associated with greater error. Additionally, greater error was observed when Eddy was used without MP-PCA denoising; error was lower in Eddy than SHORELine when denoising was used, but higher when no denoising was performed.

Unsurprisingly, in most cases, higher motion in the input data was associated with greater error following head motion correction. However, this effect was sometimes impacted by an interaction with the denoising and motion correction methods chosen. While denoising with MP-PCA had minimal impact on the error present in SHORELine output, it had a major impact on Eddy: error was systematically lower across both shelled schemes when the data was denoised first. Somewhat surprisingly, and in contrast to nearly all other parameter combinations evaluated, the amount of rotation in the input data was not associated with greater error when Eddy was used in conjunction with MP-PCA (Note: the same was not true for translations).

Error also varied substantially across acquisition schemes. For example, among shelled schemes, error was systematically lower in HCP than ABCD data. In general, acquisition schemes that sampled a greater number of directions tended to have less error following head motion correction (**Figure 4**). The interaction in **Supplementary Table 1** is driven by the exception to this trend: CS-DSI has the fewest number of directions but retained a low RMSE. As SHORELine is the only existing algorithm that can process non-shelled schemes, data from these acquisition schemes could not be evaluated using Eddy.

**Figure 4:**
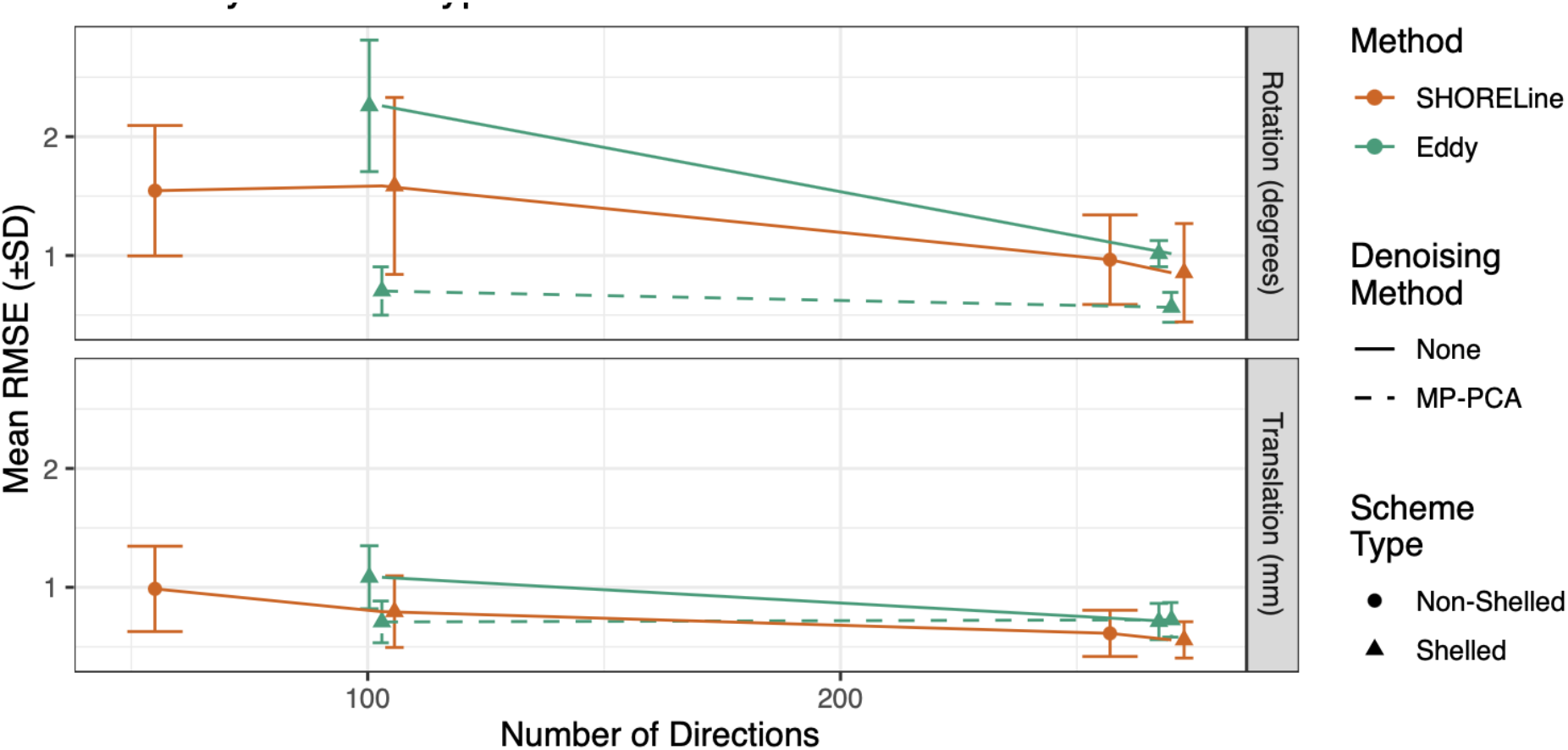
Head motion correction error varies by number of directions in sample scheme. RMSE as a function of the number of sampled directions. Points near x=103 are from the ABCD sequence; those near 270 are from the HCP; those near x=258 are from the DSIQ5; and those near x=55 are from the HASC55. SHORELine’s performance was equivalent with and without denoising, so results without denoising are shown. In general, better performance was seen with more directions and when MP-PCA was used in conjunction with Eddy.

Next, we directly compared Eddy and SHORELine (**Figure 5**) using data from the shelled ABCD and HCP sampling schemes where both methods were applicable. Overall, differences between the methods were quite small and depended in part on the use of denoising, the sampling scheme, and whether rotations or translations were evaluated (see full statistical results in **Supplementary Table 2**). For ABCD, SHORELine had less error than Eddy in all scenarios when no denoising was applied first. However, when the data was first denoised with MP-PCA, Eddy showed a slight superiority that scaled with the prevalence of motion in the input data.

**Figure 5:**
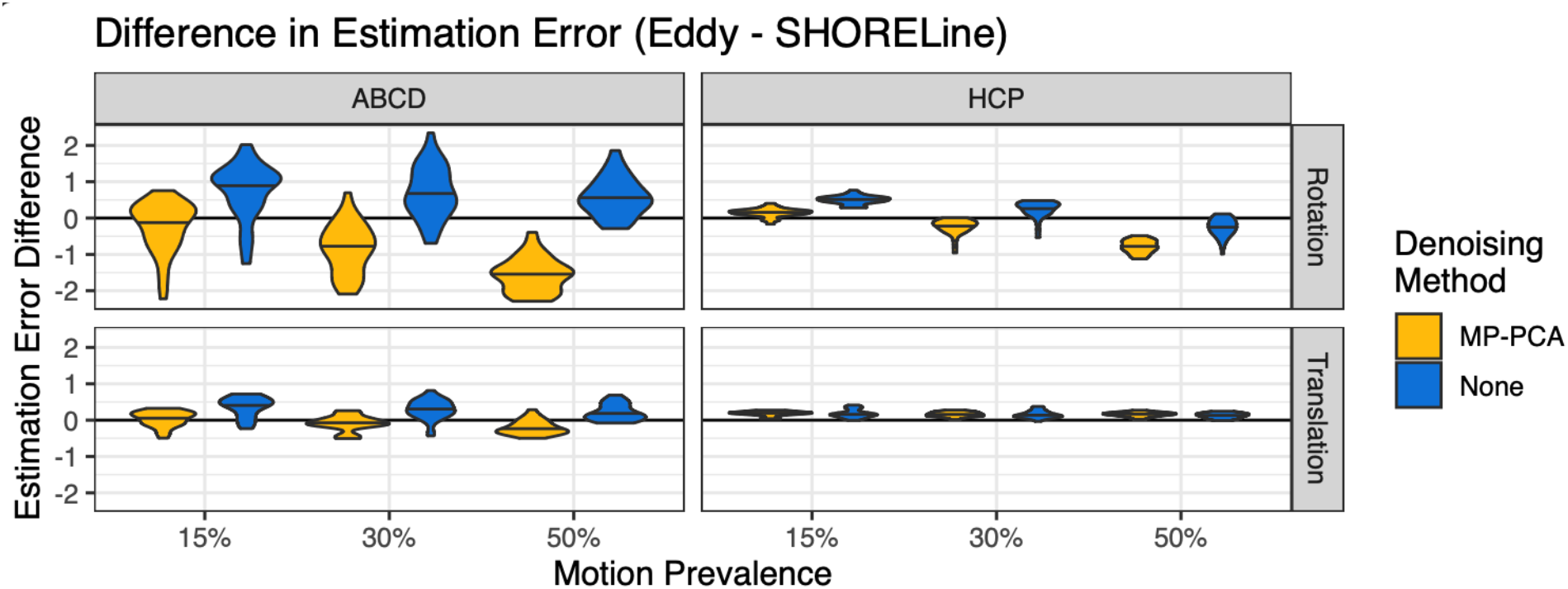
De-noising impacts relative performance of motion correction methods. Violin plots of the difference between Eddy’s and SHORELine’s RMSE for shelled schemes. Positive values indicate that SHORELine had lower RMSE than Eddy, while negative values mean Eddy had lower RMSE than SHORELine. In general, Eddy performed slightly better than SHORELine when MP-PCA denoising was performed first, whereas SHORELine was slightly superior when Eddy was used without denoising. However, overall absolute differences were quite small.

In contrast, results from the data simulated with the HCP scheme were more heterogeneous, and related to both the measure evaluated (rotation vs. translation) and the amount of motion present in the input data. For rotations, Eddy following MP-PCA outperformed SHORELine as more motion is present in the simulation. However, without MP-PCA denoising, SHORELine modestly outperformed Eddy overall for HCP data. For translations, SHORELine error was lower across all conditions, but differences were quite small.

### Output image smoothness is impacted primarily by motion present in data

Next, we evaluated the image smoothness (quantified as FWHM) of the preprocessed data following head motion correction (**Figure 6**). Image smoothness is a measure of blurring during preprocessing, which reduces anatomical detail. Ideally, preprocessing minimizes the introduction of additional image smoothness. As expected, we found that the largest driver of output image smoothness was the motion prevalence in the simulated data: across all sampling schemes and motion correction methods, more motion in the input data was associated with greater smoothness in the output images. Somewhat surprisingly, denoising did not significantly impact output image smoothness (see **Supplementary Table 3**). Additionally, although differences were small (i.e., <0.5mm FWHM), SHORELine produced significantly sharper output than Eddy.

**Figure 6:**
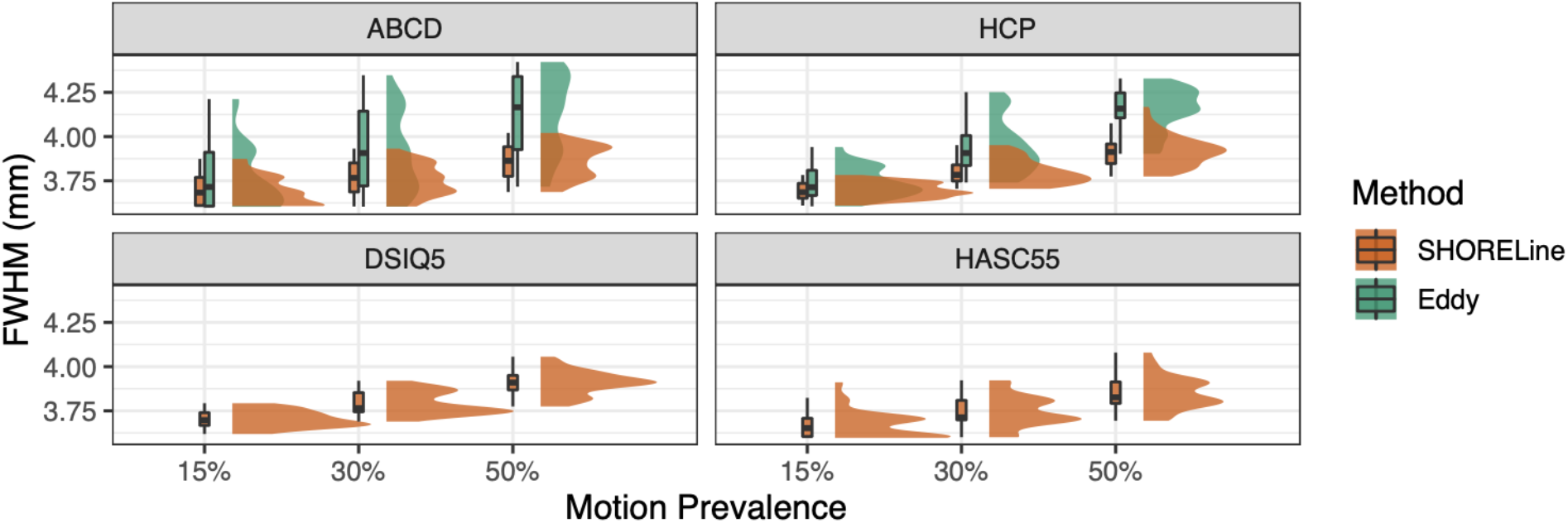
Spatial smoothness following motion correction is impacted by the amount of motion present in the input data. Both methods produce blurrier images as the amount of motion present increases. Denoising with MP-PCA did not significantly impact smoothness, so variation in de-noising is not shown. Eddy cannot process non-shelled schemes, so only data from SHORELine is shown. In general, Eddy produced blurrier images than SHORELine, although differences were small.

### Output image quality is improved by head motion correction and denoising

As a final step, we quantified the quality of the output images using the neighboring DWI correlation (NDC; **Figure 7**). We calculated NDC for both the unprocessed input data and the output from both SHORELine and Eddy; this approach allowed us to examine how much pre-processing improved data quality compared to the raw input data. Notably, head motion correction yielded substantial improvements in NDC across levels of input data motion and sampling schemes. Indeed, the improvement in NDC with preprocessing in general scaled with the prevalence of motion in the input data (see bottom row, **Figure 7**). However, in nearly all cases, greater prevalence of motion was associated with reduced output data quality even following preprocessing. One important exception to this association was the use of Eddy with HCP data, where higher prevalence of motion was not associated with reduced NDC.

**Figure 7:**
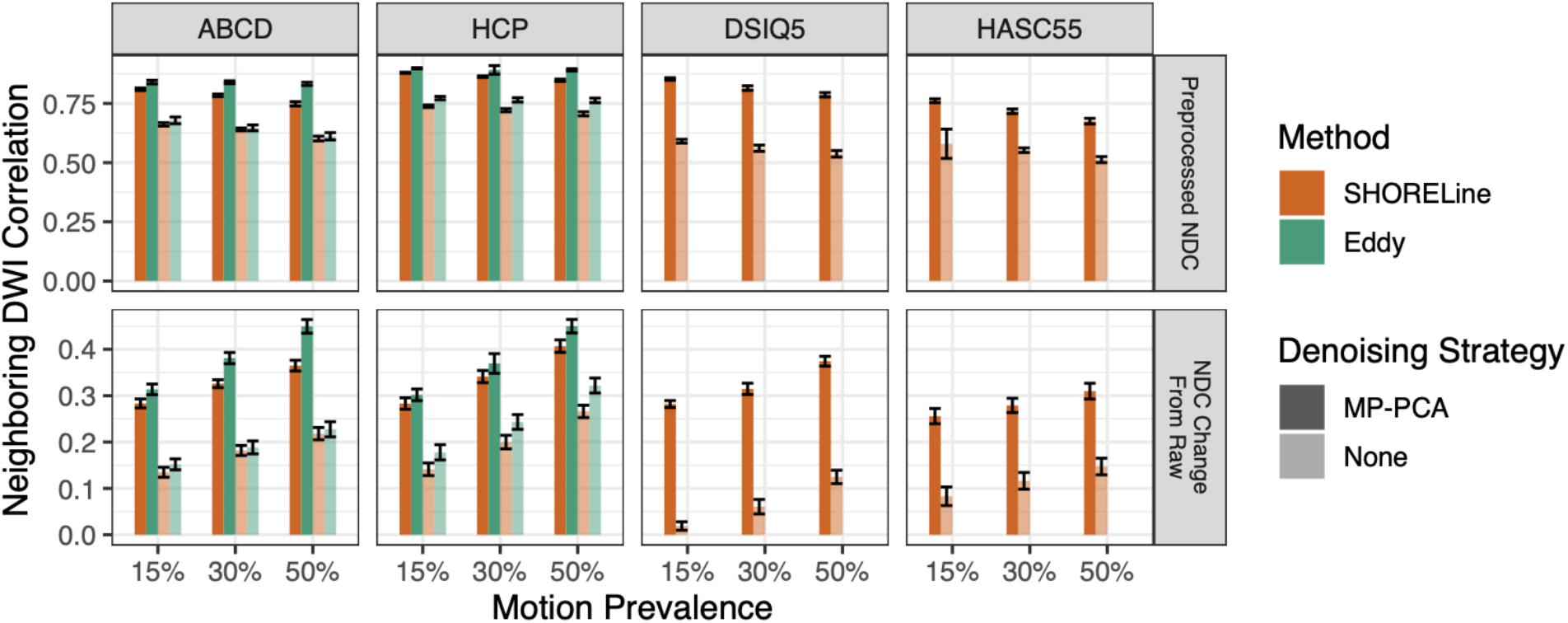
Motion correction and denoising improve data quality. Data quality was quantified as the neighboring DWI correlation (NDC). The top row displays the mean and standard deviation of the NDC values after preprocessing. The bottom row shows the change in NDC from the NDC calculated on the unprocessed scans. In general, NDC was improved by motion correction, especially following MP-PCA denoising. Eddy yielded improvements in NDC that were significantly higher than SHORELine, although differences were small.

Additionally, we found that denoising with MP-PCA improved NDC values in all scenarios that we evaluated (**Supplementary Table 4**). Denoising related improvements were particularly marked for non-shelled schemes, which could only be processed with SHORELine. Furthermore, we found that Eddy had slightly higher NDC scores than SHORELine; however, the difference was reduced when MP-PCA was not performed. The small relative advantage for Eddy over SHORELine scaled with greater motion prevalence in the input data.

## DISCUSSION

The primary finding from this large-scale evaluation is that existing tools for dMRI motion correction work—and work well. Across all methods, denoising, motion prevalence, and sampling schemes, both Eddy and SHORELine estimated head motion correctly, with observed errors being unbiased and quite small. This analysis is a necessary prerequisite to show that head motion correction is accurate on real dMRI data: there would be little hope that these methods would work on real world data if these analyses showed biased estimation of motion, or if errors of more than a fraction of a voxel translation or degree rotation were present. This outcome was clearly not the case, which should reassure users of both methods, whether they are already in wide use (Eddy) or recently introduced (SHORELine).

Although observed errors were on average quite small, our evaluation did identify several factors that influenced error magnitude. These factors included input data quality (specifically motion prevalence), denoising, and acquisition scheme. Unsurprisingly, across all scenarios examined, the single biggest determinant of the amount of error observed was the prevalence of motion in the simulated input data. Similarly, more prevalent motion in the input data resulted in greater image blurring (e.g., FWHM) and reduced data quality (as quantified by the NDC). These results emphasize that acquiring high quality data remains of critical importance for all studies, as no amount of image processing can fully compensate for extensive in-scanner motion.

Nonetheless, one preprocessing step that is not always used in standard pipelines— denoising—had remarkably beneficial effects on the outcomes we evaluated. Across all scenarios, data denoised with MP-PCA improved in quality without impacting smoothness, meaning that a simple increase in smoothness did not drive the increased performance (Woods, Grafton, Holmes, et al., 1998a; Woods, Grafton, Watson, et al., 1998b). However, the use of denoising did interact with choice of head motion correction method in an unexpected manner: while denoising did not impact the error estimated by SHORELine, estimation error was markedly reduced when Eddy was applied to data that had been first denoised by MP-PCA. This result was unanticipated: Eddy developers do not recommend denoising prior to head motion correction because of theoretical concerns regarding the way denoising might change the noise distribution in the data. When paired with MP-PCA, Eddy performed uniquely well in some contexts, with rotation error failing to scale with severity of motion. While some field-standard pipelines—such as QSIPrep – do by default apply MP-PCA denoising prior to head motion correction, other widely used pipelines (such as HCP pipelines) do not. These empirical results suggest that this recommendation may require re-evaluation, in particular for the many large-scale data resources that rely on processing pipelines that apply Eddy without denoising.

Our study considered four different acquisition schemes—including two commonly used shelled schemes and two non-shelled schemes—that allowed us to examine how outcomes were impacted by this important experimental design choice. We found that sequences with more directions tended to have lower error, likely due to the fact that increased data volume facilitated model training and fitting. However, perhaps the most important result of comparing these four schemes was the finding that non-shelled schemes could be successfully corrected as well as shelled schemes. Notably, processing of non-shelled schemes was only possible with SHORELine. As DSI methods have important advantages for modeling the average ensemble propagator, this represents a milestone for the preprocessing of non-shelled schemes and may accelerate their adoption by the neuroimaging community. Furthermore, the particularly impressive performance on the very-brief 55-direction CS-DSI scheme emphasizes the promise of compressed sensing methods for the many translational applications where scan time is limited and motion may be prominent (i.e., children and clinical populations).

In contrast to the impact of input data quality, denoising, and acquisition scheme, direct comparisons of Eddy and SHORELine were notable mainly for the small effects that were observed. There were small but significant differences in estimation error between the two methods, but the direction of the effect largely depended on whether MP-PCA was also used. As noted above, adding denoising resulted in a notable reduction in the observed error for Eddy. Images processed by SHORELine were slightly sharper than Eddy, but the magnitude of difference was of uncertain practical significance (i.e., 0.06 mm FWHM). Conversely, we found that Eddy had a higher NDC than SHORELine. However, this difference was also of unclear practical impact. To put the NDC difference in context, Eddy on average had an NDC that was 0.02 units higher than SHORELine-processed images; the difference in NDC between unprocessed data and processed images was approximately 20 times larger. Overall, our results emphasize that both methods perform quite well.

Several limitations of this study should be noted. First, we compared Eddy and SHORELine, but other dMRI head motion estimation methods exist. However, the MAPMRI-based method implemented in TORTOISE (Irfanoglu et al., 2017) is likely to perform similarly to SHORELine due to its similar method for estimating registration targets. Second, SHORELine requires sampling schemes with at least two unique non-zero *b*-values, preventing a comparison of performance on common single shell schemes. Third, real world data may differ in both the types of artifacts and the types of movement observed. For example, the FiberFox simulations do not simulate within-volume motion and slice dropout and Eddy current distortion is always linear. However, our use of simulated data allowed us to have a known ground truth by which to benchmark these methods and systematically manipulate multiple distinct parameters in a factorial design.

Moving forward, we anticipate that preprocessing methods for DWI will continue to advance. In particular, several important features that are included in Eddy—such as eddy current correction—could be included in SHORELine in future releases. Furthermore, the advent of cutting-edge denoising techniques—such as Patch2Self (Fadnavis et al., 2020)—that leverage self-supervised learning may have important implications for head motion correction. Finally, we have released all simulated images, processing software, analytic code, and results associated with this work; these may prove useful for future benchmarking efforts and facilitate comparisons to existing methods. This open and fully-reproducible workflow both bolsters confidence in the current results and may accelerate evaluation studies moving forward.

## ACKNOWLEDGMENTS

This study was supported by grants from the National Institutes of Health: R01MH112847, R01MH120482, R37MH125829, R01EB022573, R01MH113550, R01MH123550, RF1MH116920, U01AG052943, RF1MH121867, R01MH119185, R01MH120174. Additional support was provided by the AE Foundation and the Penn/CHOP Lifespan Brain Institute. STG was supported by the Institute for Collaborative Biotechnologies under Cooperative Agreement W911NF□19□2□0026 from the Army Research Office.

## SUPPLEMENTARY MATERIAL

**Supplementary Table 1.** Statistical comparisons of denoising, scheme type, number of directions, denoising method, and motion prevalence on RMSE of estimated motion.

| Predicting RMSE of Estimated Motion |  |  |  |  |  |  |
| --- | --- | --- | --- | --- | --- | --- |
| Characteristic | Translation |  |  | Rotation |  |  |
|  | Beta | 95% CI<sup>†</sup> | p-value | Beta | 95% CI<sup>†</sup> | p-value |
| (Intercept) | 0.528 | 0.487, 0.569 | <0.001 | 1.125 | 1.019, 1.231 | <0.001 |
| Scheme Type |  |  |  |  |  |  |
| *Shelled* | — | — |  | — | — |  |
| *Non-Shelled* | 0.115 | 0.068, 0.161 | <0.001 | -0.278 | -0.398, -0.157 | <0.001 |
| N. Directions | -0.001 | -0.001, -0.001 | <0.001 | -0.004 | -0.005, -0.004 | <0.001 |
| Denoising |  |  |  |  |  |  |
| *MP-PCA* | — | — |  | — | — |  |
| *None* | 0.060 | 0.039, 0.081 | <0.001 | 0.335 | 0.280, 0.391 | <0.001 |
| Motion Prevalence | 0.013 | 0.012, 0.014 | <0.001 | 0.021 | 0.019, 0.023 | <0.001 |
| Scheme Type * N. Directions |  |  |  |  |  |  |
| *Non-Shelled \* N. Directions* | -0.001 | -0.001, 0.000 | <0.001 | 0.001 | 0.001, 0.002 | <0.001 |

<sup>†</sup> CI = Confidence Interval

**Supplementary Table 2.**
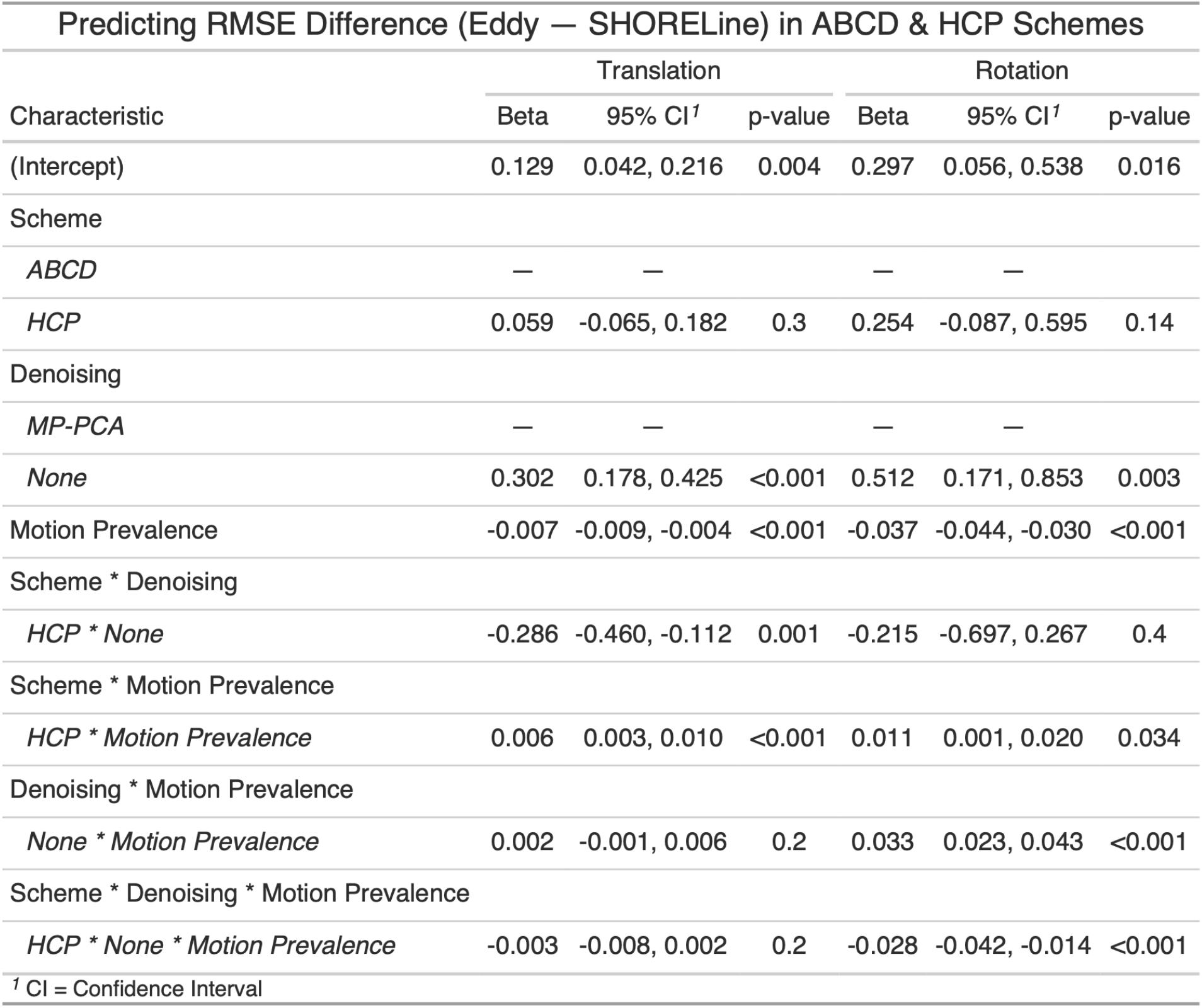
Statistical comparison of Eddy and Shoreline in shelled schemes. The outcome measure was the difference in error (RMSE) between Eddy and Shoreline (Eddy-Shoreline); the model included main effects of denoising, motion prevalence in the input data, and shell scheme; interactions of these main effects were modeled as well.

**Supplementary Table 3.** Statistical comparisons of image smoothness between Eddy and Shoreline. The outcome measure was the difference in image smoothness (FWHM) between Eddy and Shoreline (Eddy-Shoreline); the model included main effects of denoising, motion prevalence in the input data, and sampling scheme.

| Predicting FWHM (Eddy — Shoreline) of Each Scan |  |  |  |
| --- | --- | --- | --- |
| Main Effects of Percent Motion, Scheme, and Denoising |  |  |  |
| Characteristic | Beta | 95% CI<sup>†</sup> | p-value |
| (Intercept) | -0.060 | -0.083, -0.036 | <0.001 |
| Motion Prevalence |  |  |  |
| 15% | — | — |  |
| 30% | -0.086 | -0.111, -0.060 | <0.001 |
| 50% | -0.199 | -0.225, -0.174 | <0.001 |
| Scheme |  |  |  |
| ABCD | — | — |  |
| HCP | 0.018 | -0.002, 0.039 | 0.084 |
| Denoising |  |  |  |
| MP-PCA | — | — |  |
| None | -0.009 | -0.030, 0.012 | 0.4 |
| <sup>†</sup> CI = Confidence Interval |  |  |  |

**Supplementary Table 4.**
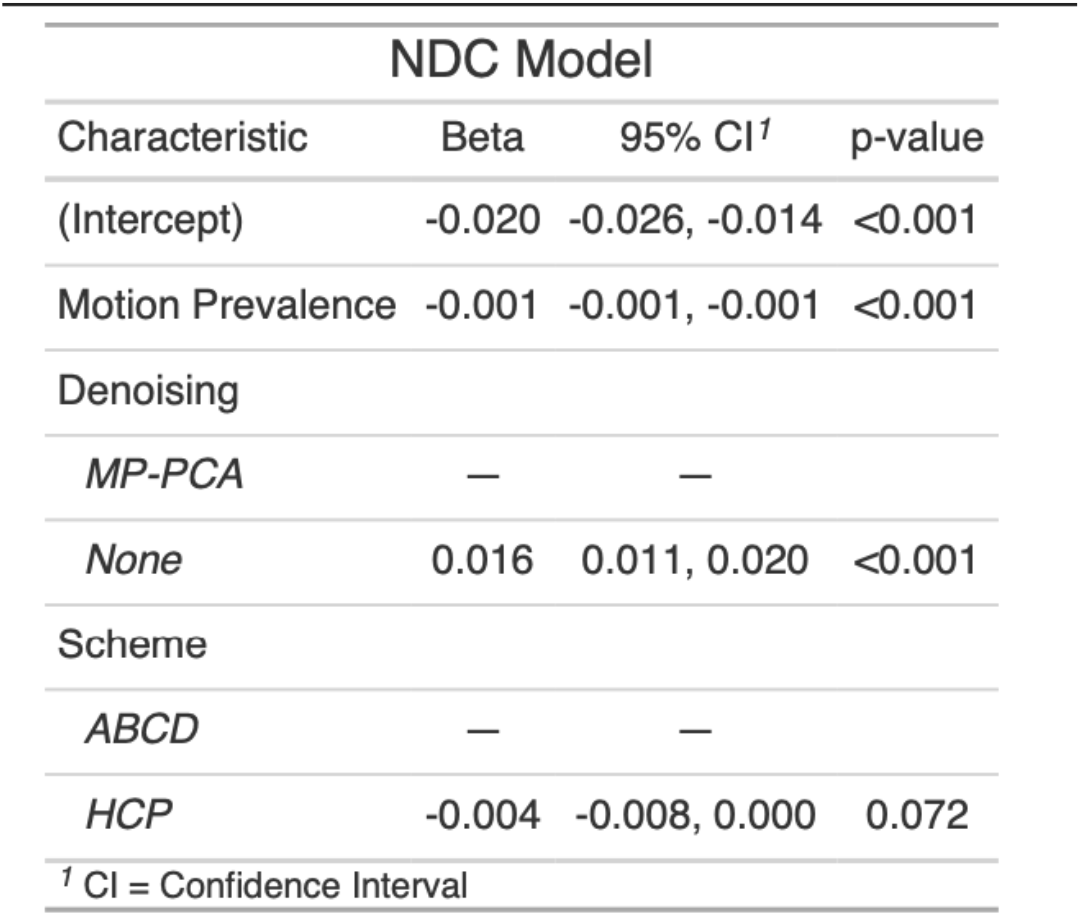
Statistical comparisons of image quality between Eddy and Shoreline. The outcome measure was the difference in image quality (NDC) between Eddy and SHORELine (Eddy-Shoreline); the model included main effects of denoising, motion prevalence in the input data, and shell scheme.

